# Locus coeruleus–hippocampal pathway activation promotes spatial memory updating during reversal learning

**DOI:** 10.64898/2026.08.26.747270

**Authors:** Coline Portet, Jyotika Bahuguna, Romain Goutagny

**Author notes:** **Corresponding authors:** Jyotika Bahuguna  & Romain Goutagny. equal contribution (Jyotika Bahuguna, Romain Goutagny).

## Abstract

Spatial navigation requires animals to integrate current environmental information with previously acquired spatial memories. The locus coeruleus provides neuromodulatory input to the hippocampus, but whether this pathway facilitates spatial learning in general or preferentially supports the updating of established representations remains unclear. Here, we selectively activated LC projections to the dorsal hippocampus while mice performed object-location recognition and an appetitive radial-maze task involving initial spatial learning followed by reversal. LC–hippocampal activation enhanced object-location memory and improved reversal learning, reducing total and working-memory errors, but did not affect initial spatial reference acquisition or retention. To characterize navigation beyond classical performance measures, we developed a graph-based analysis comparing each observed trajectory with paths generated from random, regular, small-world and heuristic goal-directed network models. Radial-maze trajectories contained a structured mixture of goal-directed-like and regular or serial-like patterns that evolved across learning. In addition, agreement with the goal-directed model was associated with fewer errors and greater proximity to the rewarded arm. Together, these findings indicate that LC inputs to the hippocampus preferentially facilitate spatial memory updating rather than uniformly enhancing spatial learning, and introduce a complementary framework for quantifying the organization of radial-maze trajectories.

## INTRODUCTION

Spatial navigation requires animals to combine information about their current environment with previously acquired spatial memories, a process that critically depends on the hippocampus [1–4]. Within the hippocampal circuit, CA1 integrates direct entorhinal inputs carrying information about the current environment with internally processed information arriving from CA3. This organization places CA1 in a position to compare ongoing sensory information with stored representations and to contribute to the updating of spatial memories when environmental conditions change [5–8]. Accordingly, hippocampal spatial representations reorganize following changes in context, task demands or reward location [9,10]. Although the hippocampal mechanisms supporting spatial coding have been extensively studied [11,12], less is known about the mechanisms which determine when existing spatial representations should be maintained and when they should be updated.

The noradrenergic system is a strong candidate for regulating this balance. The Locus Coeruleus (LC provides the main noradrenergic input to the hippocampus and modulates neuronal excitability, synaptic plasticity, encoding and memory persistence [13–15]. LC terminals can also contribute to dopamine release in the dorsal hippocampus, providing an additional mechanism through which this pathway may regulate hippocampal plasticity [16–18]. Consistent with this role, selective activation of LC projections to the dorsal hippocampus enhances several forms of hippocampus-dependent memory, including spatial, contextual and associative memory [17–20].

LC activity may be particularly important when stored representations must be modified. LC neurons respond to novelty, unexpected events and changes in behavioral contingencies, and their phasic activity can precede adaptive changes in behavior [21–23]. Phasic LC activation can induce hippocampal remapping even when the external environment remains unchanged [24], while LC input to CA1 contributes to place-cell reorganization following a change in reward location [25]. LC projections to dorsal CA1 are also required for the linking of memories acquired close in time and for updating spatial contextual recognition memory [26,27]. Together, these findings suggest that LC inputs do not simply strengthen hippocampal memories, but may facilitate the incorporation of new information into existing representations. However, it remains unclear whether activation of the LC–hippocampal pathway preferentially supports memory updating or facilitates spatial learning more generally. Previous studies have usually examined initial memory formation, contextual updating or changes in hippocampal representations in separate experimental paradigms. A direct comparison between initial spatial learning and the subsequent updating of the same learned goal representation is therefore lacking.

To address this question, we selectively activated LC projections to the dorsal hippocampus while mice performed two hippocampus-dependent spatial tasks. We first assessed object-location recognition and then used an appetitive eight-arm radial-maze task in which mice initially learned a stable reward location before the reward was moved to the opposite arm. This design allowed us to compare the effects of LC– hippocampal activation on initial spatial reference learning and reversal learning within the same behavioral framework.

Classical measures such as latency and number of errors provide essential indices of task performance, but they do not fully describe how animals organize their successive choices to reach the goal. Rodents can use allocentric information based on external spatial cues, egocentric responses, serial exploration of adjacent arms or combinations of these strategies [28,29]. In addition, the strategy expressed within a single trajectory may change as the animal explores the maze. Recent trajectory-based methods have therefore attempted to move beyond global performance measures by classifying search patterns or segmenting trajectories into distinct behavioral strategies [30]. Most of these approaches nevertheless assign trajectories or trajectory segments to discrete categories. We developed a graph-based framework that instead compares each observed arm-choice sequence continuously with trajectories generated from four reference network models. These models represented random exploration, regular or serial exploration, small-world-like exploration and goal-directed-like navigation. This approach allows an individual trajectory to express graded similarity to several navigation patterns rather than being assigned to a single strategy. Because no canonical graph representation of goal-directed navigation exists, the goal-directed network was defined as a heuristic model in which transition probabilities were concentrated along efficient routes toward the rewarded arm. We then tested whether this framework captured learning-dependent changes in trajectory organization and whether agreement with the different models was related to established measures of spatial performance.

Using this combined approach, we show that activation of LC–hippocampal projections enhances object-location memory and improves performance during reversal learning, while having no detectable effect on the initial acquisition of spatial reference memory. The graph-based analysis further shows that radial-maze trajectories contain a mixture of goal-directed-like and regular or serial-like structure and that agreement with the goal-directed model is associated with more efficient navigation. These findings support a preferential role for LC inputs to the hippocampus in spatial memory updating and introduce a complementary framework for quantifying the organization of radial-maze trajectories.

## RESULTS

### Selective targeting of LC projections to the dorsal hippocampus

To selectively activate locus coeruleus projections to the dorsal hippocampus, we combined retrograde Cre expression in the dorsal hippocampus with Cre-dependent expression of either ChETA-eYFP or eYFP in the LC. Optical fibers were implanted bilaterally above the dorsal hippocampus, allowing stimulation of LC terminals during behavioral testing. Histological analysis confirmed selective expression within the LC. Among eYFP-positive neurons, 87.4 ± 2.15% were also positive for tyrosine hydroxylase, indicating that the targeted population was predominantly noradrenergic (Figure 1a). We next examined the distribution of hippocampus-projecting neurons within the LC. Labelled neurons were more abundant in the dorsal than in the ventral LC, as indicated by a significant effect of dorsoventral position (F_(1,4)_ = 37.35, p = 0.004, η²p = 0.903; Figure 1b). There was no significant main effect of anteroposterior level (F_(3,12)_ = 2.34, p = 0.125, η²p = 0.369), although the interaction between anteroposterior level and dorsoventral position approached significance (F_(3,12)_ = 3.12, p = 0.066, η²p = 0.438). These results confirm the selective targeting of a predominantly dorsal population of LC neurons projecting to the dorsal hippocampus.

**Figure 1:**
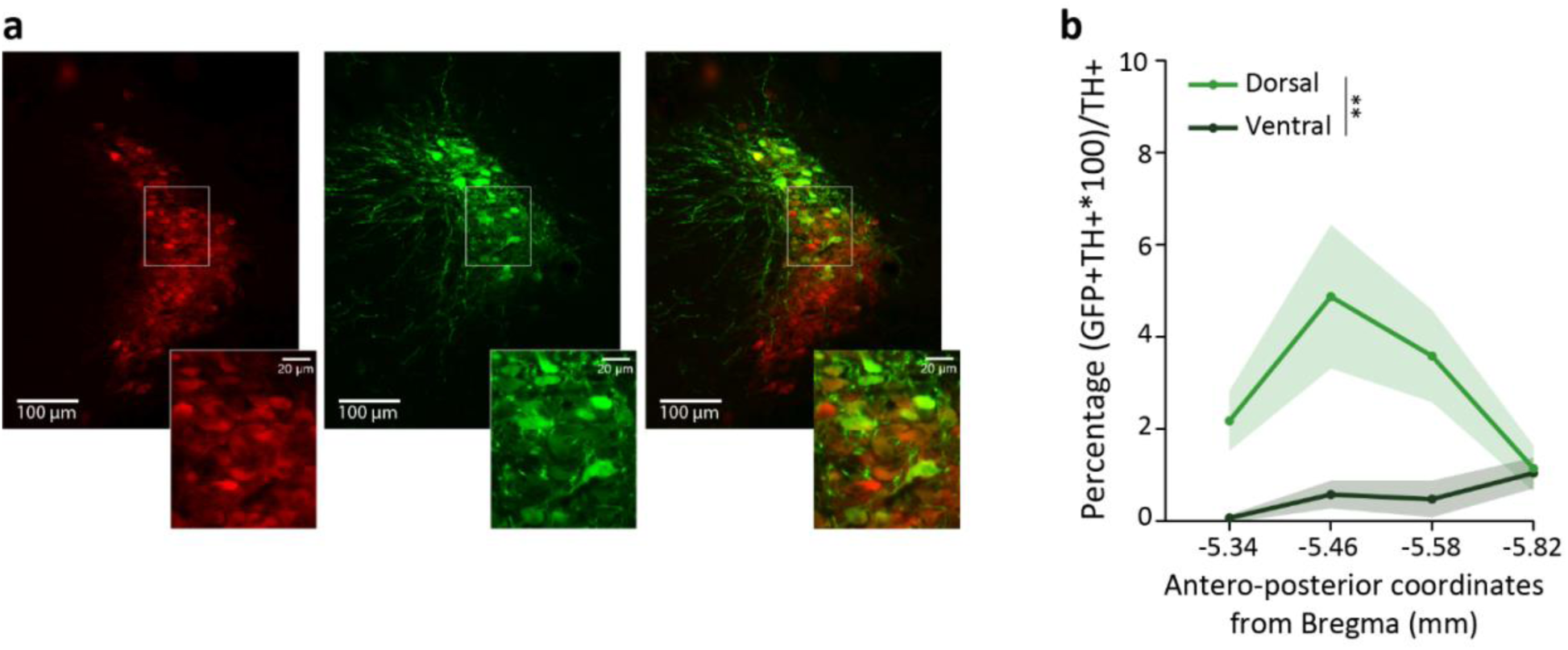
Histological control of viral infection in LC neurons. **a.** Specificity of viral infection demonstrated by immunofluorescence. From left to right: TH staining alone (red), GFP staining alone (green), and TH+ GFP+ co-staining. Scale bar represents 100 µm (whole LC image) or 20 µm (cell zoom). **b.** Distribution of LC neurons projecting to CA1 along the dorso-ventral and antero-posterior axes. Significant differences were observed between dorsal and ventral regions (global effect: p = 0.004).

### Activation of LC–hippocampal projections enhances object-location memory

We first examined whether activation of LC projections to the dorsal hippocampus enhanced hippocampus-dependent spatial memory using an object-location recognition task (Figure 2a). Analysis of the recognition index revealed significant effects of task phase (F_(1,14)_ = 7.18, p = 0.018, η²p = 0.339) and virus group (F_(1,14)_ = 6.90, p = 0.020, η²p = 0.330), together with a significant phase × group interaction (F_(1,14)_ = 5.26, p = 0.038, η²p = 0.273; Figure 2c). In ChETA mice, the recognition index increased between acquisition and test (p = 0.021), whereas no corresponding change was observed in eYFP controls (p = 1.000). Consistent with this result, ChETA mice preferentially explored the displaced object during the test phase (t_(7)_ = 3.627, p = 0.004, d = 1.28), whereas eYFP mice performed at chance level (t_(7)_ = −1.057, p = 0.836, d = −0.372). This improvement was not explained by a change in overall exploratory activity. Total object exploration time was not affected by task phase (F_(1,14)_ = 3.688, p = 0.075, η²p = 0.209), virus group (F_(1,14)_ = 0.013, p = 0.910, η²p = 0.001), or their interaction (F_(1,14)_ = 0.037, p = 0.850, η²p = 0.003; Figure 2b). Activation of LC–hippocampal projections therefore enhanced memory for object location without altering object exploration.

**Figure 2:**
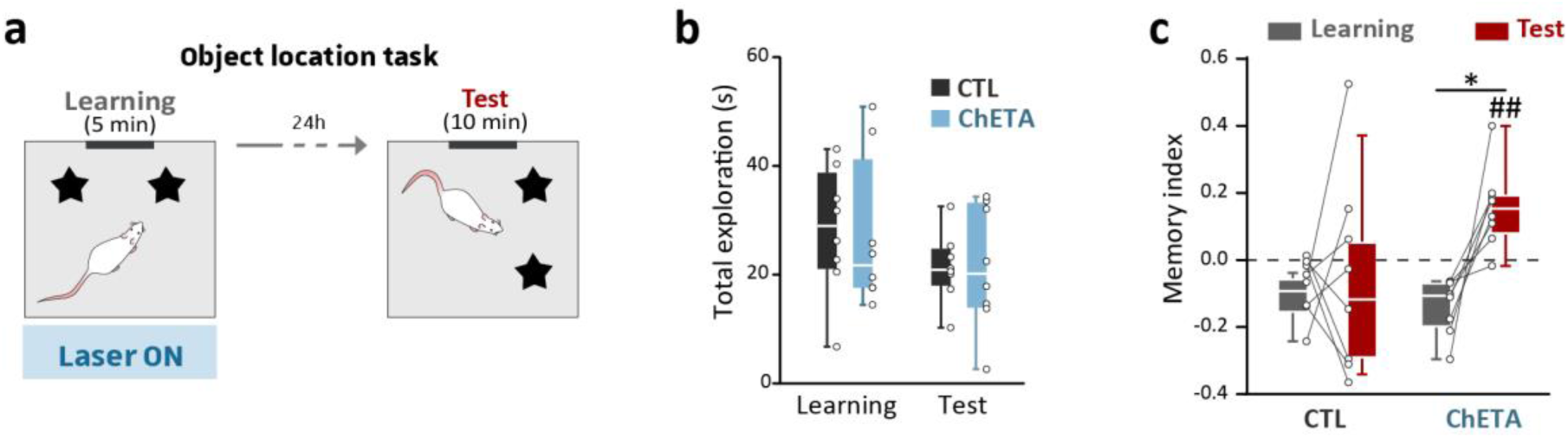
LC-Hippocampal pathway stimulation enhances performance in the object location task. **a.** Total object exploration time during sampling and test phases shows comparable exploration levels between ChETA and eYFP groups. **b.** Recognition index for the displaced object during acquisition and recall phases. An index greater than zero indicates preferential exploration of the displaced object compared to the fixed object. During the test phase, only ChETA mice showed significant discrimination of the displaced object (## : p = 0.004), also significantly different from the memory index from the learning phase (* : p = 0.021).

### LC–hippocampal activation does not broadly enhance spatial learning

The object-location experiment showed that activation of LC–hippocampal projections could facilitate spatial memory. We next asked whether this effect reflected a general enhancement of spatial learning or depended on the demands placed on previously acquired spatial representations. Mice were trained for ten days to locate a rewarded arm in an eight-arm radial maze. The rewarded location remained stable relative to extra-maze cues throughout this initial learning phase (Figure 3a). Both ChETA and eYFP mice progressively improved their performance. Training day significantly affected total errors (F_(9,72)_ = 3.821, p < 0.001, η²p = 0.323), spatial errors (F_(9,72)_ = 7.627, p < 0.001, η²p = 0.488), working-memory errors (F_(9,72)_ = 4.153, p < 0.001, η²p = 0.342), and latency to reach the rewarded arm (F_(9,72)_ = 18.185, p < 0.001, η²p = 0.694; Figure 3b). Despite this robust learning, LC–hippocampal activation did not improve initial acquisition. There was no significant main effect of virus group on total errors (F_(1,8)_ = 0.429, p = 0.531, η²p = 0.051), spatial errors (F_(1,8)_ = 0.571, p = 0.472, η²p = 0.067), working-memory errors (F_(1,8)_ = 1.40, p = 0.270, η²p = 0.149), or latency (F_(1,8)_ = 0.0356, p = 0.855, η²p = 0.004). Similarly, no virus group × training day interaction was observed for total errors (F_(9,72)_ = 0.403, p = 0.930, η²p = 0.048), spatial errors (F_(9,72)_ = 0.381, p = 0.847, η²p = 0.062), working-memory errors (F_(9,72)_ = 0.385, p = 0.871, η²p = 0.059), or latency (F_(9,72)_ = 0.746, p = 0.666, η²p = 0.085). During the subsequent probe test, both groups spent more time than expected by chance in the target arm (ChETA: t_(5)_ = 3.445, p = 0.009, d = 1.406; eYFP: t_(3)_ = 7.879, p = 0.002, d = 3.940; Figure 3c). Target-arm preference did not differ between groups (t_(8)_ = −0.786, p = 0.454, d = −0.507). Thus, both groups learned and remembered the initial reward location, with no detectable effect of LC–hippocampal activation. The facilitatory effect observed in the object-location task therefore did not reflect a uniform enhancement of spatial learning.

**Figure 3:**
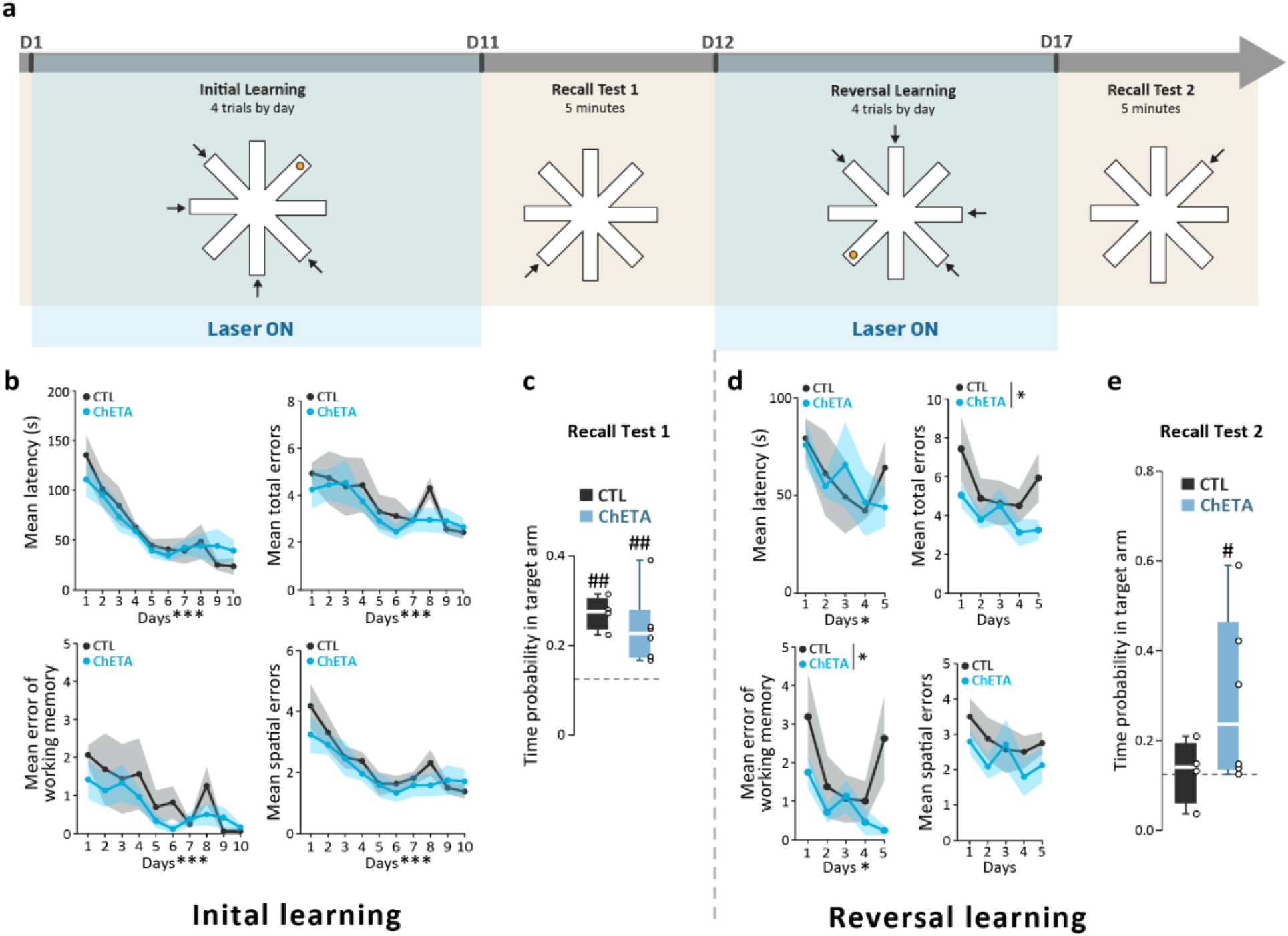
Spatial reference memory performance during learning and reversal phases of the task. **a.** Schematic of the spatial reference memory task, testing both initial learning and reversal in the eight-arm maze. LC–hippocampal pathway activation (blue areas) occurred only during the trial. Arrows indicate the possible starting arms relative to the position of the rewarded arm (yellow circle). **b and d.** Evolution of mean latency, total errors, working-memory errors, and spatial errors across days for ChETA and eYFP mice during initial learning (b) and reversal learning (d). **c and e**. Relative time spent in the target arm during the probe test following initial learning (c) and the probe test following reversal (e), relative to chance level (dashed line). Both ChETA and eYFP groups performed above chance during the initial probe test; during the reversal probe test, only the ChETA group performed above chance. Comparisons to chance level are indicated by # (p < 0.05) or ## (p < 0.01); comparisons between groups are indicated by * (p ≤ 0.05), ** (p ≤ 0.01), and *** (p ≤ 0.001).

### LC–hippocampal activation facilitates adaptation to a new reward location

We next asked whether LC–hippocampal activation facilitated adaptation when the reward location changed. After initial learning, the rewarded arm was moved to the opposite side of the maze, and mice were trained for five additional days. Performance continued to evolve during reversal. Training day significantly affected working-memory errors (F_(4,32)_ = 2.760, p = 0.044, η²p = 0.257) and latency to reach the rewarded arm (F_(4,32)_ = 2.679, p = 0.049, η²p = 0.251; Figure 3d). The effects of day on total errors (F_(4,32)_ = 2.192, p = 0.092, η²p = 0.215) and spatial errors (F_(4,32)_ = 1.269, p = 0.303, η²p = 0.137) did not reach significance. In contrast to initial acquisition, LC–hippocampal activation improved performance during reversal. ChETA mice made fewer total errors than eYFP controls (F_(1,8)_ = 5.690, p = 0.044, η²p = 0.416) and fewer working-memory errors (F_(1,8)_ = 8.500, p = 0.019, η²p = 0.515). No significant group effect was observed for spatial errors (F_(1,8)_ = 1.310, p = 0.285, η²p = 0.141) or latency (F_(1,8)_ = 0.0128, p = 0.913, η²p = 0.002). The virus group × training day interaction was not significant for any performance measure. During the reversal probe test, ChETA mice spent more time than expected by chance in the new target arm (t_(5)_ = 2.152, p = 0.042, d = 0.879), whereas eYFP mice did not show a significant preference for this location (t_(3)_ = 0.186, p = 0.432, d = 0.093; Figure 3e). The direct comparison between groups did not reach significance (t_(8)_ = 1.590, p = 0.151, d = 1.020). Together, these results show that LC–hippocampal activation improved performance when the previously learned reward location was no longer valid. This effect was expressed as a reduction in total errors and repeated visits to previously explored arms, supporting a preferential contribution of this pathway to spatial memory updating rather than to initial spatial acquisition.

### Graph-based analysis reveals structured navigation across learning and reversal

The classical performance measures established that LC–hippocampal activation selectively improved reversal learning. However, error counts and latency provide little information about how animals organize their successive choices within individual trials. We therefore developed a graph-based analysis to characterize the structure of radial-maze trajectories beyond these global measures. Each observed trajectory was compared with paths generated from four reference network models representing random exploration, regular or serial exploration, small-world-like exploration, and goal-directed-like navigation. The goal-directed model was defined by concentrating transition probabilities along efficient routes between the starting arm and the rewarded arm.

We first quantified how this heuristic model topology differed from the canonical reference networks in terms of frequently used graph metrics (Figure S1). The goal-directed network combined sparse connectivity with short paths and high global efficiency. It also showed low transition entropy and marked heterogeneity in strengths of the incoming transition weights, reflecting the concentration of transition probabilities onto a restricted set of routes leading toward the goal. In contrast, the random network distributed transitions broadly, the regular network favored local serial transitions, and the small-world network combined local structure with occasional longer-range transitions. These differences provided a structural basis for comparing observed trajectories with distinct navigation patterns.

During initial learning, trajectory similarity differed strongly among network models (F_(3,1200)_ = 358.98, p < 0.001, η²p = 0.473; Figure 4a; Full omnibus statistics for the mixed-effects models during initial and reversal learning are reported in Table S1). Similarity also changed across training days (F_(1,390)_ = 41.35, p < 0.001, η²p = 0.096), and this evolution differed among network models, as indicated by a significant network type × day interaction (F_(3,1200)_ = 26.47, p < 0.001, η²p = 0.062). Across both groups, observed trajectories were most similar to the goal-directed model, followed by the regular model. Similarity to the random and small-world models was lower. Post hoc comparisons confirmed that similarity to the goal-directed model was higher than similarity to each of the other models in both ChETA and eYFP mice (all Bonferroni-corrected p < 0.001; Table S3). Across both groups, trajectories showed the greatest similarity to the goal-directed model, followed by the regular model, indicating that radial-maze navigation was characterized primarily by goal-directed-like and serial-like organization (Table S2).

**Figure 4.**
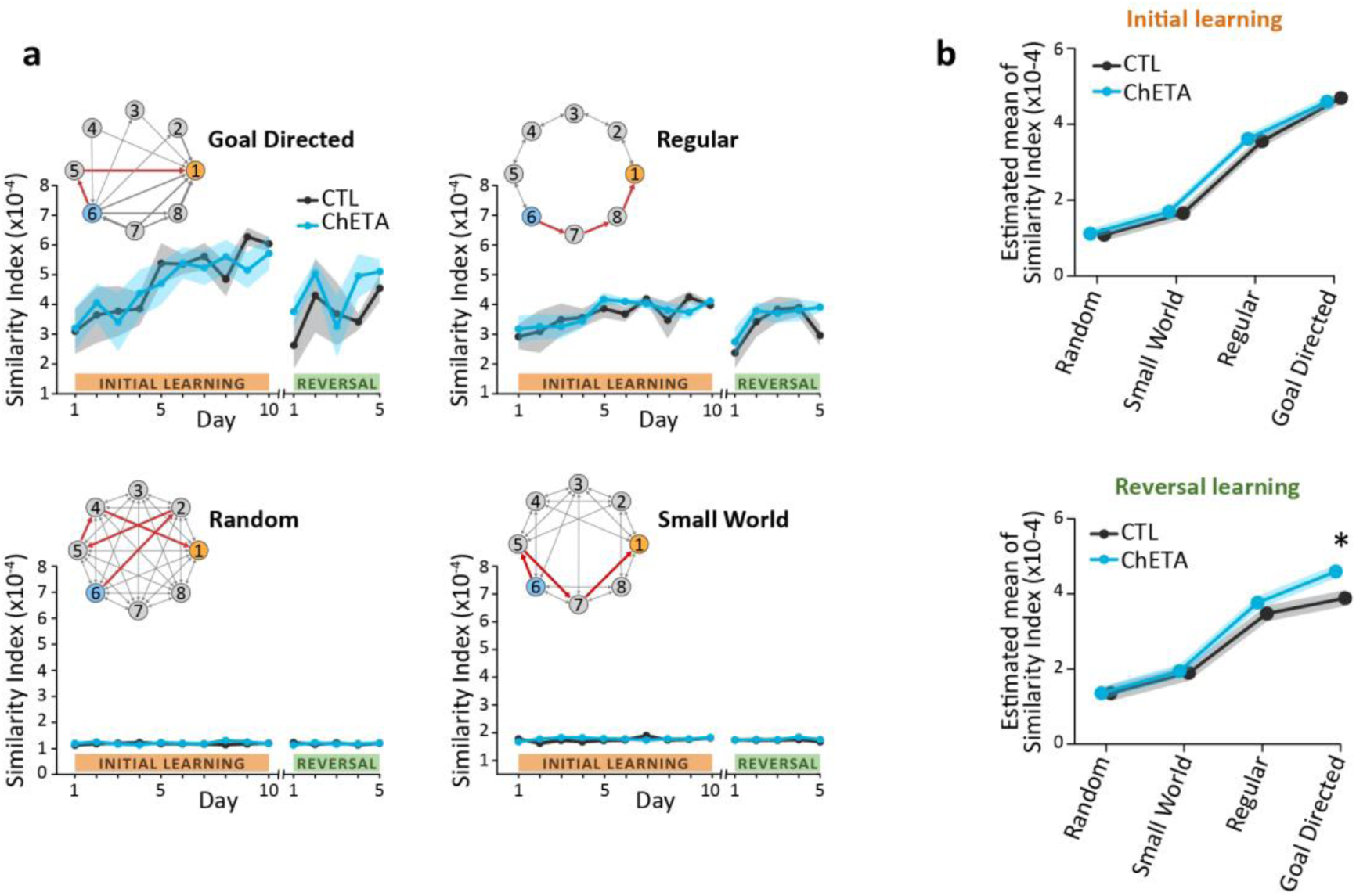
Graph-based characterization of radial-maze trajectories during initial and reversal learning. **a.** Similarity between observed mouse trajectories and trajectories generated from four reference network models representing random, small-world-like, regular or serial, and goal-directed-like navigation. For each model, nodes represent the eight maze arms and directed edges indicate possible transitions between arms, with edge thickness proportional to transition probability. Example simulated trajectories from the starting arm, shown in blue, to the rewarded arm, shown in yellow, are indicated in red. Across initial learning and reversal, observed trajectories were most similar to the goal-directed and regular models, whereas similarity to the random and small-world models was lower. **b.** Estimated marginal means of trajectory similarity for the four reference models during initial learning and reversal learning. No group difference was observed during initial acquisition. During reversal, a planned contrast within the goal-directed model showed greater goal-directed-like similarity in ChETA mice than in eYFP controls (p = 0.015), with no corresponding group differences for the random, regular, or small-world models.

Consistent with the classical performance measures, the broad organization of trajectories was comparable between groups during initial acquisition. There was no significant effect of virus group (F_(1,11)_ = 0.03, p = 0.86, η²p = 0.003), virus group × network type interaction (F_(3,1200)_ = 0.11, p = 0.96, η²p = 0.001), or virus group × network type × day interaction (F_(3,1200)_ = 0.33, p = 0.81, η²p = 0.001). During reversal, trajectories remained structured rather than becoming random following reward relocation. Similarity differed among network models (F_(3,600)_ = 66.34, p < 0.001, η²p = 0.249), with goal-directed-like and regular-like patterns remaining predominant. A significant effect of day further indicated continued reorganization of trajectories during reversal learning (F_(1,190)_ = 4.18, p = 0.042, η²p = 0.022). The network type × day interaction approached significance (F_(3,600)_ = 2.45, p = 0.063, η²p = 0.012). Although the overall virus group × network type interaction was not significant (F_(3,600)_ = 0.56, p = 0.645, η²p = 0.003), a planned contrast within the goal-directed model showed greater goal-directed-like similarity in ChETA mice than in eYFP controls during reversal (t_(61)_ = −2.50, p = 0.015, d = -0.403; Table S4). No corresponding group differences were observed for the random, regular, or small-world models. To determine whether the graph-defined patterns were related to behavioral efficiency, we calculated the likelihood of each observed transition sequence under the four reference models using a measure independent of the trajectory-similarity score (Figure S2; see Methods *“Log-Likelihood of observed trajectories*”). Agreement with the goal-directed model closely tracked performance across initial learning and reversal. Higher goal-directed likelihood was associated with fewer working-memory errors in both ChETA and eYFP mice (r = −0.62 ± 0.20 and r = −0.75 ± 0.20, respectively) and with greater proximity to the rewarded arm (r = 0.86 ± 0.05 and r = 0.82 ± 0.09, respectively). These relationships were weaker for the random, regular and small-world models. The graph-based analysis therefore identified predominantly goal-directed-like with a mixture of serial-like trajectory organization that evolved across learning. During reversal, LC–hippocampal activation was associated with a selective increase in goal-directed-like similarity, while the broader organization across network models remained unchanged. By preserving the sequence of arm choices within each trial, this framework captured a behaviorally meaningful dimension of navigation that was not available from latency or error counts alone.

## DISCUSSION

This study examined whether locus coeruleus inputs to the hippocampus facilitate spatial learning in general or preferentially support the updating of previously acquired spatial information. Activation of LC projections to the dorsal hippocampus enhanced object-location memory and improved performance when the rewarded location was changed, but did not affect the initial acquisition or retention of spatial reference memory. Together, these results suggest that LC–hippocampal activation is more effective during spatial memory updating than during initial learning. The graph-based analysis provided a complementary description of this behavior, showing that radial-maze trajectories contained a structured mixture of goal-directed-like and regular or serial-like patterns that evolved across learning.

The improvement in object-location memory confirms previous evidence that LC inputs to the dorsal hippocampus facilitate hippocampus-dependent memory [17–20]. LC terminals can release both noradrenaline and dopamine in the hippocampus, and both neuromodulators regulate neuronal excitability, synaptic plasticity and memory formation [14–18]. The present experiment does not distinguish their respective contributions. However, the absence of any change in total object exploration indicates that the improvement cannot be readily explained by increased exploratory activity or a nonspecific effect of stimulation on behavioral engagement. In contrast, LC–hippocampal activation did not improve the initial acquisition of the radial-maze task. Both groups progressively reduced their errors and latency and subsequently showed a preference for the rewarded arm during the probe test. This absence of facilitation is important because it argues against a uniform enhancement of spatial learning. It differs from the improvement in Barnes-maze performance reported after activation of LC terminals in the dorsal hippocampus [17]. Several differences between the tasks may contribute to this discrepancy. The Barnes maze is generally based on escape from an exposed environment, whereas the present radial-maze task used an appetitive reward and repeated training. LC recruitment is sensitive to arousal, novelty, uncertainty and behavioral relevance, and its influence may therefore depend on the motivational context and difficulty of the task [21–23]. In addition, the extended acquisition protocol may have allowed control mice to reach a high level of performance, leaving limited room for further improvement. Whatever the underlying explanation, our results show that stimulation of this pathway is not sufficient to improve spatial performance under all learning conditions.

A clear benefit emerged when the rewarded arm was moved to the opposite side of the maze. During reversal, ChETA mice made fewer total errors and fewer repeated visits to previously explored arms than controls. Only ChETA mice also spent more time than expected by chance in the new target arm during the subsequent probe test, although the direct comparison between groups was not significant. The reduction in errors during training therefore provides the strongest evidence that LC–hippocampal activation facilitated adaptation to the new reward location. Importantly, this effect was observed without a corresponding reduction in latency. Stimulation did not simply make the animals reach the end of the trial more rapidly. Instead, it improved the efficiency with which they sampled the maze after the spatial contingency changed. This preferential effect during reversal is consistent with an emerging view of the LC as a regulator of memory updating. LC neurons respond to unexpected changes and can rapidly modify their responses when the behavioral significance of a stimulus changes [22,23]. Within the hippocampus, phasic LC activation can induce remapping in an otherwise familiar environment [24], and LC input to CA1 contributes to the reorganization of place-cell activity following reward relocation [25]. LC projections to dorsal CA1 are also required for updating spatial contextual recognition memory, an effect that depends particularly on dopaminergic signaling through D1/D5 receptors [27]. Moreover, direct β-adrenergic stimulation of the dorsal dentate gyrus has been reported to improve reversal learning while disrupting retrieval when no update is required, supporting the idea that noradrenergic signaling can favour a transition from retrieval toward encoding of new information [24]. Together with these studies, our results suggest that LC input may promote the reconfiguration of hippocampal representations when current information conflicts with a previously established memory. This interpretation does not imply that LC–hippocampal activity is exclusively involved in updating. Activation of this pathway can facilitate initial encoding under other experimental conditions [17,19,20]. Rather, its influence may be greatest when novelty, prediction error or changed task contingencies indicate that an existing representation is no longer sufficient. In this framework, catecholaminergic input would alter the balance between maintaining a stable hippocampal representation and allowing it to be modified. The present study supports this view by comparing initial learning and reversal within the same animals, using the same maze, reward and stimulation protocol. The behavioral dissociation therefore cannot be attributed solely to differences between experimental apparatuses or pathway manipulations.

The graph-based analysis extended the behavioral characterization beyond conventional error and latency measures. Radial-maze trajectories were most similar to the goal-directed model, but they also contained a substantial regular or serial-like component. This suggests that mice did not rely on a single, fixed navigation pattern. Instead, their trajectories combined efficient movements toward the goal with structured transitions between neighbouring arms. Importantly, these patterns changed across acquisition and continued to evolve during reversal, showing that the analysis was sensitive to the reorganization of behavior with learning. During reversal, ChETA mice also showed greater similarity to the goal-directed model in a planned within-model contrast. This effect was modest and occurred in the absence of a significant virus group × network type interaction, suggesting a selective increase in goal-directed-like trajectory organization rather than a broader reconfiguration across navigation models. The behavioral relevance of the graph framework was further supported by the transition-likelihood analysis. Trajectories that were more probable under the goal-directed model were strongly correlated with the temporal dynamics of both behavioral measures: tracking the decrease in working-memory errors during learning, their transient increase during reversal, and subsequent decline, as well as the reciprocal pattern of proximity to the rewarded arm, which increased with learning, decreased during reversal, and increased again thereafter. These relationships are partly expected because the model was constructed around efficient routes to the goal. Nevertheless, their consistency across animals and task phases shows that the model captured a dimension of trajectory organization related to successful navigation. The graph-based approach therefore complements classical performance measures by preserving the sequence of choices made within a trial. It also allows each trajectory to express similarity to several reference patterns, rather than assigning the complete path to a single categorical strategy. The reference networks should not be interpreted as direct estimates of the cognitive processes used by the animals. In particular, similarity to the goal-directed model does not demonstrate an exclusively allocentric strategy. The model describes trajectories biased toward efficient routes to the rewarded arm, regardless of the information used by the animal to generate those routes. For this reason, the term goal-directed-like is more appropriate than allocentric. The framework should instead be considered a quantitative description of trajectory structure that can generate hypotheses about navigation strategies. Its generality will need to be tested in independent datasets, other maze geometries and experiments in which allocentric, egocentric and serial strategies can be dissociated through specific manipulations.

Several other limitations should be considered. The number of mice completing the radial-maze experiment was modest, particularly in the control group. Although the effects on total and working-memory errors during reversal were accompanied by large effect sizes, their magnitude should be confirmed in a larger cohort. In addition, only male mice were included, and whether these findings generalize to females remains to be established. The present experiments also do not identify the cellular mechanisms responsible for the behavioral improvement. LC terminals in the hippocampus can engage both noradrenergic and dopaminergic receptors, and the relative contribution of these systems may depend on the behavioral condition and hippocampal subregion [16–18,27]. Combining projection-specific stimulation with receptor-selective pharmacology or direct measurements of catecholamine release will be necessary to determine how these signals contribute to spatial memory updating.

In conclusion, activation of LC projections to the dorsal hippocampus enhanced object-location memory and improved adaptation to a relocated reward, while leaving initial spatial reference learning unchanged. These findings support a preferential contribution of LC–hippocampal inputs when established spatial information must be revised in response to changing contingencies. The graph-based analysis further revealed that radial-maze navigation is organized through a combination of goal-directed-like and regular or serial-like trajectory patterns that evolve with learning. Together, the results identify a selective role for the LC–hippocampal pathway in spatial memory updating and provide a complementary framework for describing how animals organize their choices during spatial navigation.

## Acknowledgements

We acknowledge the In Vitro Imaging Platform – Strasbourg (CNRS UAR3156), member of the national infrastructure France-BioImaging supported by the French National Research Agency (ANR-10-INBS-04).

## Author contributions

CP performed the experiments and analyzed the data; JB conceptualized and implemented the graph network analysis; RG conceived the experimental and analytical design; all authors wrote the article

## Data availability

The datasets generated and analyzed during the current study are available from the corresponding authors upon reasonable request. The code used for the graph-based trajectory analyses will be made publicly available through the FunSy team GitHub repository: (https://github.com/FunSyCNRS).

## METHODS

### Animals

Male CD1 mice (n = 18), obtained from Charles River Laboratories France (Saint-Germain-Nuelles, France), were 8 weeks old at the start of the experiments and were individually housed in Makrolon cages (16 × 32 × 14 cm) with bedding and nesting material. Housing conditions were strictly controlled (24±1°C, 30±5% humidity, 12h light/dark cycle with lights on at 8:00, background noise 45±5dB). Animals had ad libitum access to food and water except during behavioral testing periods requiring water restriction. All experimental protocols agreed with the European Committee Council directive (2010/63/UE) and were approved by the French Ministry of Research (APAFIS#20388-2019042517013497). All animal were used for behavioral tasks, and only a subset (n = 5) was used for anatomical characterization of the projecting LC neurons to the dorsal hippocampus.

### Surgical Procedures

Animals were anesthetized with isoflurane (4% induction, 0.5-2% maintenance) and placed in a stereotaxic frame. Body temperature was maintained using a heating pad with rectal probe feedback. Local anesthesia (Lurocaine 2mg/kg and Bupivacaine 2mg/kg, s.c.) and systemic analgesia (Metacam 1mg/kg) were administered before surgery. The skull was exposed and aligned to ensure flat-skull configuration.

A retrograde AAV expressing Cre recombinase (pENN.AAV.hSyn.Cre.WPRE.hGH) was injected bilaterally into dorsal hippocampus (0.8µL per site at 100nL/min; coordinates from bregma: AP: -2.46mm, ML: ±1.75mm, DV: -1.5mm from dura), followed by a 10-minute waiting period. A Cre-dependent AAV expressing either ChETA-eYFP (pAAV-Ef1a-DIO-ChETA-eYFP; experimental group, n = 8) or eYFP alone (pAAV-EF1a-DIO-eYFP-WPRE-HGHpA; control group, n = 8) was injected bilaterally into LC (0.4µL per site at 100nL/min; AP: -5.4mm, ML: ±0.9mm, DV: -3.75mm).

Optical fibers (200µm diameter) were implanted bilaterally above dorsal hippocampus (AP: -2.46mm, ML: ±2.25mm, DV: -1.42mm). Implants were secured using UV-curable adhesive followed by dental cement (Paladur, Kulzer).

### Behavioral Testing

All behavioral experiments began 3 weeks after surgery. Experimenters were blind to experimental groups.

#### Object Location Recognition

Testing occurred in a black plexiglass arena (55×55x40cm) with a visual cue (A4-sized black/white striped card) on one wall. After three days of habituation (10min daily), mice explored two identical objects for 5min (acquisition). Twenty-four hours later, the least explored object during acquisition was displaced to a new spatial location and exploration was assessed for 5min (test). Object exploration was scored when the mouse’s nose was within 2cm of an object. Recognition index was calculated as a ratio with a chance level at 0 using the formula:

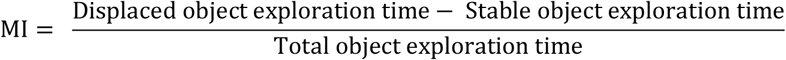

Two animals were excluded after sampling phase due to an absence of object exploration; the resulting experimental group included 16 animals (ChETA n = 8; eYFP n = 8).

#### Spatial Reference Memory (ATA task)

The ATA task was performed as previously described (Douchamps et al., 2024). Briefly, testing was conducted in an elevated (65cm) eight-arm radial maze with 55cm long arms radiating from a 52cm diameter central platform. To ensure sufficient motivation, mice were placed on water restriction with 1h daily access to water (∼1h30 after the end of the session). Animals were first habituated to maze components by exploring a single isolated arm baited with water reward (100μL, 5% sucrose) for four consecutive days. Subsequently, mice were familiarized with the complete apparatus over two days, with all arms baited and free exploration allowed for 10 minutes. The learning phase consisted of ten days of training with four daily trials. During each trial (maximum duration 3min), mice had to locate a single rewarded arm, with the reward location remaining constant relative to extra-maze cues. To promote allocentric navigation strategies, the starting arm varied between trials according to a pseudo-random sequence. Each trial ended either upon reward consumption or at the 3min time limit. Between trials, mice were returned to their home cage for a 5min inter-trial interval, during which the maze was cleaned with 35% ethanol to eliminate olfactory cues. Twenty-four hours after the final training day, a probe test was conducted where mice explored the maze for 5min with no reward present. Five minutes after this probe trial, a standard rewarded trial was performed to prevent extinction learning. The following day, a behavioral flexibility phase began, consisting of five days of reversal learning where the reward was relocated to the opposite arm. This phase followed the same daily protocol as the initial learning phase and concluded with a probe test conducted under identical conditions. Performance was measured by the average daily number of errors (visit to wrong arms) during learning and the relative time in target arm during probe tests (with a chance level at 0.125). In addition, we computed the strategy used by the animals to solve the maze: a trial was classified as spatial when the animal made its first visit either directly to the target arm or to one of its adjacent arms immediately followed by a visit to the target arm. Finally, we also computed working memory errors defined as the number of visits to previously explored arms. Eight animals were excluded from the task before its end due to a decrease of motivation to explore the maze (no exploration for 8 consecutive trials); the resulting experimental group is composed of 10 animals (ChETA n = 6; eYFP n = 4).

### Optogenetic Stimulation

Light delivery (473nm, 20Hz, 5s every 15s) was controlled by a pulse generator (A-M Systems) and maintained at 10mW output power, verified before each session. During behavioral testing, animals were connected to optical patch cables but stimulation was delivered only during specified acquisition/learning phases.

### Immunohistochemistry

To validate viral expression and analyze the distribution of LC neurons projecting to hippocampus, mice were deeply anesthetized an intraperitoneal injection of ketamine (200 mg/kg) and xylazine (30 mg/kg, i.p.) and transcardially perfused with phosphate-buffered saline (PBS 0.1M, pH 7.4, 4°C) containing heparin (0.1%), followed by 4% paraformaldehyde (PFA). Brains were removed, post-fixed in 4% PFA at 4°C for 24h, and cryoprotected in 20% sucrose at 4°C for 48h. Brains were then rapidly frozen in isopentane (-35 to -45°C, 1 min) and stored at -80°C until sectioning.

Coronal sections (30 µm) were collected on gelatin-coated slides using a cryostat (Leica HM500) at -20°C. Sections containing hippocampus (bregma -1.3 to -3.51 mm) and LC (bregma -4.71 to -6.11 mm) were stored at -20°C until immunostaining. For double immunofluorescence labeling, slides were first rinsed three times in PBS (0.01 M) for 10 minutes each. Non-specific binding was blocked using 5% horse serum in PBS for 1h at room temperature. Primary antibodies against tyrosine hydroxylase (TH-F11, mouse monoclonal, 1:50, Santa Cruz Biotechnology) and GFP (rabbit polyclonal, 1:200, Molecular Probes) were diluted in PBS containing 2% horse serum, 0.02% Merthiolate, and 0.5% Triton X-100. Sections were incubated with primary antibodies for 72h at 4°C with gentle agitation (250 rpm).

Following primary antibody incubation, sections were thoroughly rinsed in ultrapure water and coverslipped using DAPI Fluoromount mounting medium to visualize cell nuclei. Images were acquired using a Nanozoomer S60 (Hamamatsu) and analyzed using QuPath software (v0.2.3). To quantify viral specificity, we calculated the percentage of eYFP-expressing neurons that co-labeled with TH. The topographical distribution of hippocampus-projecting LC neurons was analyzed across four standardized anteroposterior levels (bregma -5.34, -5.46, -5.58, and -5.82 mm). At each level, the LC was divided into dorsal and ventral regions relative to its centroid, and the percentage of TH+ neurons expressing eYFP was quantified in each region. Cell counting was performed by experimenters blind to experimental conditions.

For anatomical analysis, we used a minimum of 4 sections per anteroposterior level per animal (n = 5). Neurons were considered positively labeled if their signal intensity exceeded background by at least two standard deviations. Co-labeling was confirmed by examining optical sections through the entire thickness of each cell. The total numbers of TH+ and eYFP+ neurons, as well as double-labeled neurons, were counted in each region to determine the specificity of viral expression and analyze the topographical organization of hippocampus-projecting LC neurons.

### Graph metrics

All graph analyses were performed on weighted, directed graphs using the Brain Connectivity Toolbox and NetworkX in Python.

For each network, we quantified six metrics for random, small-world, regular and goal-directed networks for 100 realizations of the graph.

Network density (D): defined as the fraction of possible directed edges present in the graph. For a directed graph with N nodes and E edges:

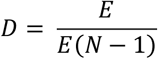

Characteristic path length (L): defined as the mean shortest-path distance between all connected node pairs in the binary graph. Let d_ij_ denote the shortest-path distance between nodes i and j.

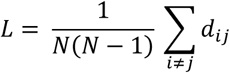

Global efficiency (E_g_): defined as the average inverse shortest-path distance between node pairs. Let d_ij_ denote the shortest-path distance between nodes i and j

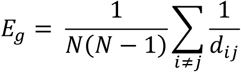

Node in-strength heterogenity (CV_in_): To quantify the heterogeneity of incoming transition strengths across nodes, we computed the coefficient of variation (CV) of node in-strength, where node in-strength was defined as the sum of incoming edge weights. For node i, w_ij_ represents the weight of the incoming edge from node j to node i, and σ(s^in^) and μ(s^in^) denote the standard deviation and mean of the weight distributions:

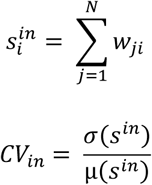

Transition entropy (H): defined as a measure of the uncertainty associated with selecting the next transition from each node. For each node i, Shannon entropy was computed from the normalized outgoing transition probabilities (p_ij_),

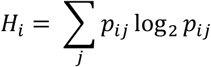

Modularity index: quantifies the extent to which the graph can be partitioned into communities with denser within-community than between-community connections and was estimated using the Louvain algorithm for weighted directed networks (*bct.modularity_louvain_dir*).

### Graph-based trajectory analysis

To quantify navigation strategies in the radial maze, we developed a graph-based trajectory analysis in which each arm of the maze was represented as a node and transitions between arms were represented as weighted edges. For each behavioral trial, the observed trajectory was defined as the ordered sequence of arms visited by the animal, starting from the initial arm and ending either at reward retrieval or at the end of the trial. This observed sequence was then compared with trajectories generated from four reference network models designed to capture different navigation patterns: random exploration, regular/serial exploration, small-world-like exploration, and goal-directed-like navigation.

The random network was used as a null model for unstructured exploration. In this model, transition probabilities between arms were uniformly distributed, so that no transition was favored over another. The regular network was designed to capture serial exploration around the maze. In this model, transitions were restricted to neighboring arms, producing trajectories in which animals move sequentially between adjacent arms. The small-world network was generated using a Watts–Strogatz-like rewiring procedure. Starting from a regular network connecting neighboring arms, edges were rewired with probability (p = 0.1), allowing mostly local transitions with occasional long-range jumps. This model was intended to capture trajectories combining local serial exploration with less frequent distant transitions.

The goal-directed network was designed to model trajectories biased toward the rewarded arm. For each trial, an all-to-all weighted graph was first generated in which edge weights reflected the physical distance between arms, with a small amount of noise added to account for uncertainty in distance estimation. The ten shortest paths between the starting arm and the rewarded arm were then identified. A new weighted graph was constructed by retaining edges that belonged to these shortest paths. Edge weights were increased in inverse proportion to the length of the path to which they belonged, such that edges contributing to shorter paths toward the goal received stronger weights. The resulting adjacency matrix was normalized to obtain transition probabilities. This model therefore captured goal-directed-like trajectories biased toward short paths leading to the rewarded arm, without assuming that animals used a purely allocentric strategy.

For each trial and each reference model, 100 network instances were generated when applicable. From each instance, 20 simulated paths were sampled using the same starting arm and the same path length as the observed trajectory. Matching the simulated and observed path lengths ensured that similarity scores were not trivially driven by the number of arms visited. The observed trajectory was then compared with the simulated paths using a weighted Jaro–Levenshtein similarity score. This score combines edit-distance-based similarity between ordered arm sequences with the probability of the path under the corresponding reference model. Higher values indicate greater similarity between the animal’s observed trajectory and the trajectory structure expected from a given network model. For each trial, this procedure yielded four similarity scores, one for each reference network model. These scores were then used as trial-level observations in the linear mixed-effects analyses described below. Because the goal-directed model captures paths biased toward the rewarded arm and its neighboring shortest routes, but does not directly prove the use of an allocentric strategy, we refer to this measure as goal-directed-like similarity throughout the manuscript.

### Log-Likelihood of observed trajectories

To provide an independent validation of the graph-based trajectory similarity analysis, we quantified the probability assigned by each reference network to the observed mouse trajectories. For each trial, the transition probability matrix of the reference network was first row-normalized so that the outgoing edge weights from each node summed to one. The log-likelihood of the observed trajectory was then computed as the mean logarithm of the transition probabilities for all consecutive arm transitions, where *v_t_* denotes the arm visited at time *t*, *L* is the trajectory length, *P(v_t_+1*∣*v_t_)* is the transition probability defined by the reference network, and *ε=10−4* was added to avoid undefined values for zero-probability transitions.

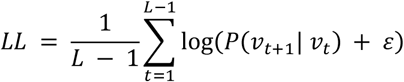

Averaging the log-likelihood across transitions yielded a measure independent of trajectory length, allowing direct comparison across trials. Higher (i.e., less negative) log-likelihood values indicate greater agreement between the observed trajectory and the corresponding navigation model.

### Working memory errors

Working-memory errors were defined as repeated visits to previously sampled non-goal arms within a trial.

### Proximity score

To quantify how closely an animal’s trajectory remained to the rewarded location, we computed a proximity score based on the graph distance between each visited arm and the goal arm. Let *d(v_t_,g)* denote the shortest-path distance between the visited arm *v_t_* and the rewarded arm *g*. The proximity score was defined as *PS*, where *L* is the number of arm visits in the trajectory. Taking the inverse of the mean distance yields larger values for trajectories that remain closer to the rewarded arm throughout the trial and smaller values for trajectories that explore arms farther from the goal.

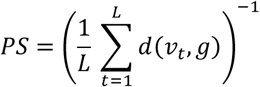

### Algorithm for generating goal-directed network

Input: (Number of maze arms: *N*, Start arm *s*, Goal arm *g*) Output: Goal-directed transition matrix *T*

- Arrange the N maze arms on a circle.
- Construct a complete directed graph G in which every pair of distinct nodes is connected.
- Assign each edge a cost equal to the Euclidean distance between the corresponding maze arms.
- Compute k shortest paths from the start node *s* to the goal node *g*
- Initialize the transition matrix *T* with zeros.
- For each shortest path *p:*

- Compute its total path length
- Set path weight w = 1/L
- For every consecutive node pair (u,v) in path *p*: T(u,v) ← T(u,v) + w
- Normalize *T* so that the sum of all edge weights equals 1

### Statistical analysis

Data are presented as mean ± SEM unless otherwise stated. Statistical analyses were performed in R and/or GraphPad Prism, with significance set at p < 0.05. All analyses were conducted by experimenters blind to experimental group. Effect sizes were reported as Cohen’s d for t-tests and partial eta-squared, η²p, for classical ANOVA models and Type III tests of fixed effect in the linear mixed model.

For the object-location task, recognition index and total exploration time were analyzed using repeated-measures two-way ANOVAs, with virus group, ChETA versus eYFP, as the between-subject factor and test phase, sampling versus recall, as the within-subject factor. When significant effects or interactions were detected, post-hoc comparisons were performed with Bonferroni correction. Recognition indices during recall were also compared with chance level, defined as 0, using one-sample t-tests.

For radial-maze acquisition and reversal learning, daily performance measures, including total errors, spatial errors, working-memory errors, and latency to reach the rewarded arm, were analyzed using repeated-measures two-way ANOVAs, with virus group as the between-subject factor and training day as the within-subject factor. Probe-test performance was assessed by comparing the relative time spent in the target arm with chance level, defined as 0.125 for the eight-arm maze, using one-sample t-tests. Direct comparisons between ChETA and eYFP mice during probe tests were performed using independent-samples t-tests. Post-hoc tests were Bonferroni-corrected when multiple comparisons were performed.

Network path similarity data were analyzed using linear mixed-effects models. The length-normalized weighted Jaro–Levenshtein similarity score was used as the dependent variable. Initial learning and reversal learning were analyzed separately. For each phase, fixed effects included network type (random, regular, small-world, goal-directed), experimental group (ChETA, eYFP), day of testing, and all interaction terms. Animal identity was included as a random intercept to account for repeated observations within animals. Because each behavioral trajectory generated four similarity scores, one for each reference network model, trial identity was also included as a random intercept in the full models to account for the non-independence of similarity scores derived from the same trajectory.

The full model for each phase was specified as:

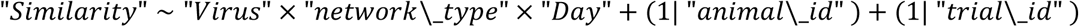

Models were fitted using maximum likelihood. Omnibus effects were assessed using type III analyses of variance with Satterthwaite approximation for denominator degrees of freedom. Estimated marginal means and post-hoc pairwise contrasts were computed using the ‘emmeans’ package (version 1.11.1), with Bonferroni correction for comparisons between network types.

