## supplementary data for "Locus coeruleus–hippocampal pathway activation promotes spatial memory updating during reversal learning"

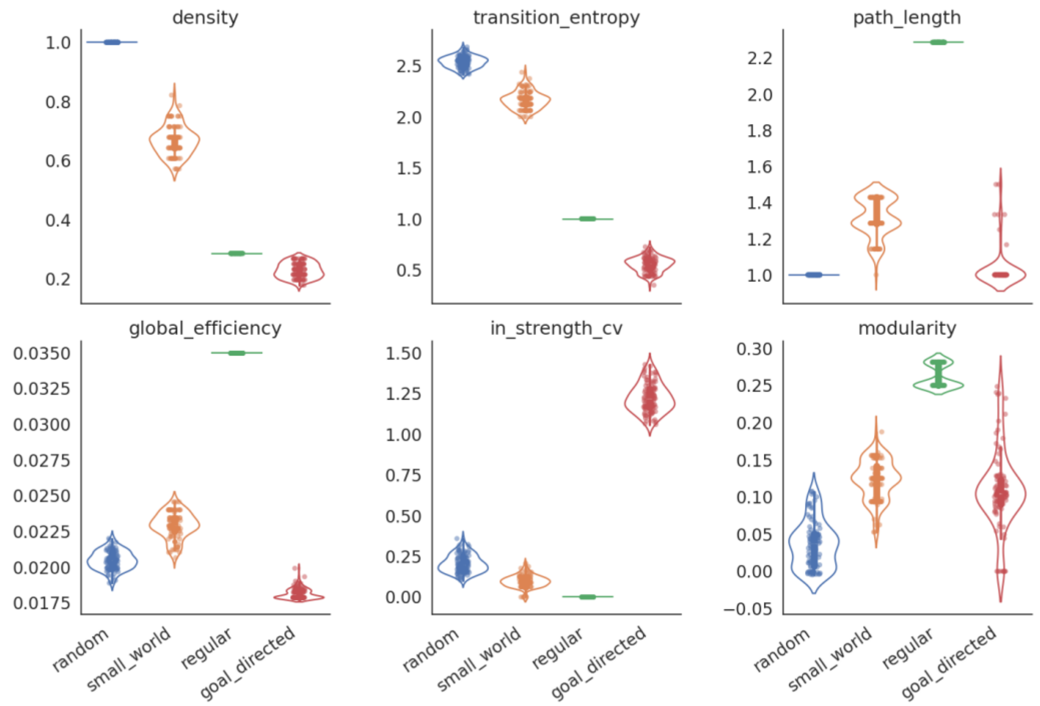


**<u>Figure S1.</u> Characterization of the proposed goal-directed network relative to canonical graph models (random, small world and regular)**.

Violin plots showing the distributions of graph metrics for ensembles of random,small-world, regular and goal-directed networks. Individual network realizations are overlaid as dots. Metrics are arranged as follows: top row, network density, transition entropy, and characteristic path length; bottom row, global efficiency, coefficient of variation (CV) of node in-strength, and modularity. Compared with the canonical graph models, the proposed goal-directed network exhibited a sparse topology with short characteristic path lengths, higher heterogeneity of incoming transition strengths, and substantially lower transition entropy, indicating that transition probabilities were concentrated onto a restricted subset of nodes forming efficient routes toward the rewarded location.

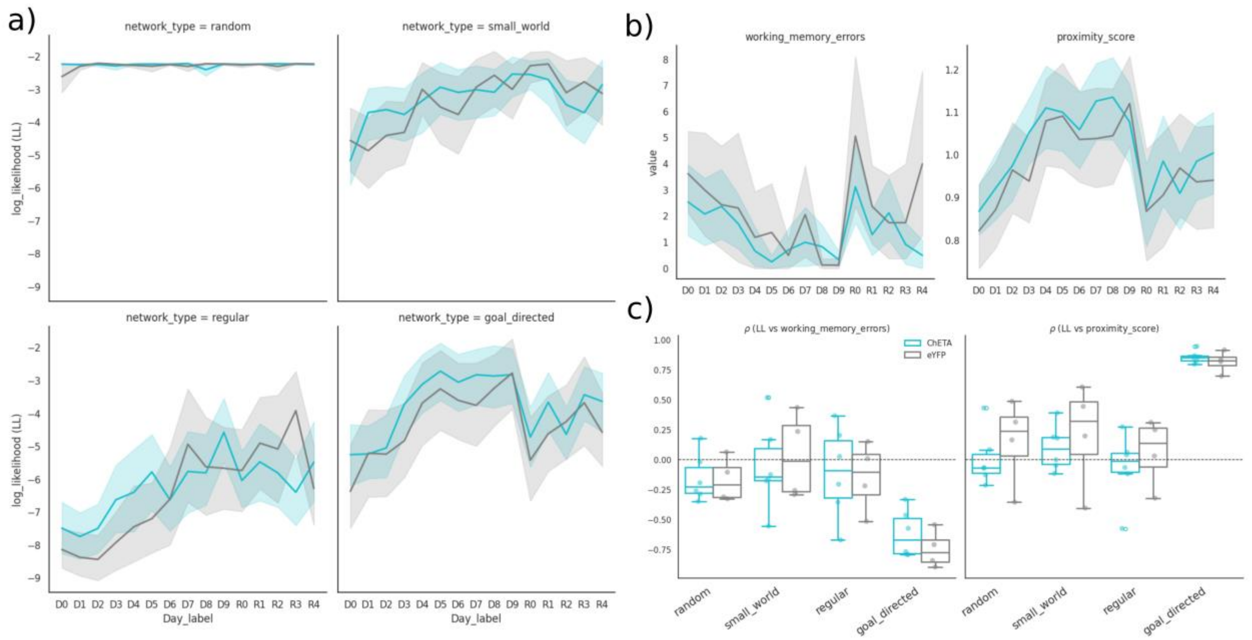


**<u>Figure S2.</u> Log-likelihood analysis provides complementary support for the goal-directed network model.**

***a.*** Mean log-likelihood of observed mouse trajectories under the four reference network models (goal-directed, regular, small-world, and random) across initial learning and reversal. Log-likelihood was computed from the row-normalized transition probability matrix of each network and represents the probability of the observed sequence of arm transitions under the corresponding navigation model. Higher (less negative) values indicate greater agreement between the observed trajectory and the network model. Owing to its dense transition matrix, the random network assigns relatively high probability to a broad range of trajectories and therefore exhibits consistently higher log-likelihood values than the other network models. In contrast, log-likelihood under the goal-directed, regular, and small-world networks changes systematically across learning and reversal, with the largest modulation observed for the goal-directed network. **b**. Working-memory errors progressively decreased during acquisition, transiently increased following reward relocation, and declined again as animals acquired the new reward location. The proximity score showed the complementary pattern, increasing during acquisition, decreasing immediately after reversal, and recovering with continued training. **c**. Pearson correlations between log-likelihood and behavioral metrics over all days (learning + reversal) for the four network types (points indicate individual mice). The goal-directed network exhibited the strongest negative correlation with working-memory errors (ChETA: −0.62 ± 0.20; eYFP: −0.75 ± 0.20) and the strongest positive correlation with proximity score (ChETA: 0.86 ± 0.05; eYFP: 0.82 ± 0.09), whereas correlations for the random, regular, and small-world networks were substantially weaker. These findings indicate that trajectories assigned higher probability by the goal-directed model were consistently associated with more efficient navigation.

**Table S1. Omnibus Type III tests for the graph-based linear mixed-effects models.**

| Phase | Effect | NumDF | DenDF | F value | P value | η²p |
|---|---|---|---|---|---|---|
| Initial learning | Virus type | 1 | 11 | 0.03 | 0.86 | 0.003 |
| Initial learning | Network type | 3 | 1200 | 358.98 | < 2e-16 | 0.473 |
| Initial learning | Day | 1 | 390 | 41.35 | 3.7e-10 | 0.096 |
| Initial learning | Virus type x network type | 3 | 1200 | 0.11 | 0.96 | 0.001 |
| Initial learning | Virus type x Day | 1 | 390 | 0.44 | 0.51 | 0.001 |
| Initial learning | Network type x Day | 3 | 1200 | 26.47 | < 2e-16 | 0.062 |
| Initial learning | Virus type x network type x Day | 3 | 1200 | 0.33 | 0.81 | 0.001 |
| Reversal learning | Virus type | 1 | 45 | 1.22 | 0.275 | 0.027 |
| Reversal learning | Network type | 3 | 600 | 66.34 | < 2e-16 | 0.249 |
| Reversal learning | Day | 1 | 190 | 4.18 | 0.042 | 0.022 |
| Reversal learning | Virus type x network type | 3 | 600 | 0.56 | 0.645 | 0.003 |
| Reversal learning | Virus type x Day | 1 | 190 | 0.03 | 0.873 | 0.001 |
| Reversal learning | Network type x Day | 3 | 600 | 2.45 | 0.063 | 0.012 |
| Reversal learning | Virus type x network type x Day | 3 | 600 | 0.06 | 0.982 | 0.001 |

**Table S2. Estimated marginal means for trajectory similarity by phase, virus type, and network model.**

| Phase | Virus type | Network model | EMM | SE | df | Lower 95% CI | Upper 95% CI |
|---|---|---|---|---|---|---|---|
| Initial learning | eYFP | Goal-directed | 0.000479 | 1.48e-05 | 40 | 0.000449 | 0.000509 |
| Initial learning | eYFP | Random | 0.000118 | 1.48e-05 | 40 | 8.77e-05 | 0.000148 |
| Initial learning | eYFP | Regular | 0.000365 | 1.48e-05 | 40 | 0.000335 | 0.000395 |
| Initial learning | eYFP | Small-world | 0.000175 | 1.48e-05 | 40 | 0.000145 | 0.000205 |
| Initial learning | ChETA | Goal-directed | 0.000469 | 1.21e-05 | 40 | 0.000444 | 0.000493 |
| Initial learning | ChETA | Random | 0.000121 | 1.21e-05 | 40 | 9.64e-05 | 0.000145 |
| Initial learning | ChETA | Regular | 0.000371 | 1.21e-05 | 40 | 0.000347 | 0.000396 |
| Initial learning | ChETA | Small-world | 0.000178 | 1.21e-05 | 40 | 0.000154 | 0.000203 |
| Reversal learning | eYFP | Goal-directed | 0.000372 | 2.19e-05 | 61 | 0.000328 | 0.000416 |
| Reversal learning | eYFP | Random | 0.000118 | 2.19e-05 | 61 | 7.46e-05 | 0.000162 |
| Reversal learning | eYFP | Regular | 0.000330 | 2.19e-05 | 61 | 0.000286 | 0.000374 |
| Reversal learning | eYFP | Small-world | 0.000173 | 2.19e-05 | 61 | 0.000129 | 0.000217 |
| Reversal learning | ChETA | Goal-directed | 0.000443 | 1.79e-05 | 61 | 0.000407 | 0.000479 |
| Reversal learning | ChETA | Random | 0.000118 | 1.79e-05 | 61 | 8.23e-05 | 0.000154 |
| Reversal learning | ChETA | Regular | 0.000359 | 1.79e-05 | 61 | 0.000324 | 0.000395 |
| Reversal learning | ChETA | Small-world | 0.000178 | 1.79e-05 | 61 | 0.000142 | 0.000213 |

**Table S3. Pairwise contrasts between network models within each virus group.**

| Phase | Virus type | Contrast | Estimate | SE | df | t ratio | Bonferroni-corrected p-value | Cohen's d |
|---|---|---|---|---|---|---|---|---|
| Initial learning | eYFP | Goal-directed - random | 3.62e-04 | 1.71e-05 | 1200 | 21.101 | < .0001 | 2.359 |
| Initial learning | eYFP | Goal-directed - regular | 1.14e-04 | 1.71e-05 | 1200 | 6.665 | < .0001 | 0.745 |
| Initial learning | eYFP | Goal-directed - small-world | 3.04e-04 | 1.71e-05 | 1200 | 17.751 | < .0001 | 1.985 |
| Initial learning | eYFP | Random - regular | -2.47e-04 | 1.71e-05 | 1200 | -14.435 | < .0001 | -1.614 |
| Initial learning | eYFP | Random - small-world | -5.74e-05 | 1.71e-05 | 1200 | -3.350 | 0.0050 | -0.374 |
| Initial learning | eYFP | Regular - small-world | 1.90e-04 | 1.71e-05 | 1200 | 11.086 | < .0001 | 1.239 |
| Initial learning | ChETA | Goal-directed - random | 3.48e-04 | 1.40e-05 | 1200 | 24.871 | < .0001 | 2.270 |
| Initial learning | ChETA | Goal-directed - regular | 9.75e-05 | 1.40e-05 | 1200 | 6.968 | < .0001 | 0.636 |
| Initial learning | ChETA | Goal-directed - small-world | 2.91e-04 | 1.40e-05 | 1200 | 20.761 | < .0001 | 1.895 |
| Initial learning | ChETA | Random - regular | -2.51e-04 | 1.40e-05 | 1200 | -17.903 | < .0001 | -1.634 |
| Initial learning | ChETA | Random - small-world | -5.75e-05 | 1.40e-05 | 1200 | -4.110 | 0.0003 | -0.375 |
| Initial learning | ChETA | Regular - small-world | 1.93e-04 | 1.40e-05 | 1200 | 13.793 | < .0001 | 1.259 |
| Reversal learning | eYFP | Goal-directed - random | 2.53e-04 | 2.78e-05 | 600 | 9.106 | < .0001 | 1.440 |
| Reversal learning | eYFP | Goal-directed - regular | 4.17e-05 | 2.78e-05 | 600 | 1.497 | 0.8096 | 0.237 |
| Reversal learning | eYFP | Goal-directed - small-world | 1.99e-04 | 2.78e-05 | 600 | 7.153 | < .0001 | 1.131 |
| Reversal learning | eYFP | Random - regular | -2.12e-04 | 2.78e-05 | 600 | -7.609 | < .0001 | -1.203 |
| Reversal learning | eYFP | Random - small-world | -5.43e-05 | 2.78e-05 | 600 | -1.953 | 0.3078 | -0.309 |
| Reversal learning | eYFP | Regular - small-world | 1.57e-04 | 2.78e-05 | 600 | 5.656 | < .0001 | 0.894 |
| Reversal learning | ChETA | Goal-directed - random | 3.25e-04 | 2.27e-05 | 600 | 14.288 | < .0001 | 1.845 |
| Reversal learning | ChETA | Goal-directed - regular | 8.34e-05 | 2.27e-05 | 600 | 3.670 | 0.0016 | 0.474 |
| Reversal learning | ChETA | Goal-directed - small-world | 2.65e-04 | 2.27e-05 | 600 | 11.671 | < .0001 | 1.507 |
| Reversal learning | ChETA | Random - regular | -2.41e-04 | 2.27e-05 | 600 | -10.618 | < .0001 | -1.371 |
| Reversal learning | ChETA | Random - small-world | -5.95e-05 | 2.27e-05 | 600 | -2.618 | 0.0544 | -0.338 |
| Reversal learning | ChETA | Regular - small-world | 1.82e-04 | 2.27e-05 | 600 | 8.000 | < .0001 | 1.033 |

**Table S4. ChETA versus eYFP contrasts within each network model.**

| Phase | Network model | EMM eYFP | EMM ChETA | Contrast | Estimate | SE | df | t ratio | P value | Cohen's d |
|---|---|---|---|---|---|---|---|---|---|---|
| Initial learning | Goal-directed | 0.000479 | 0.000469 | eYFP - ChETA | 1.04e-05 | 1.91e-05 | 40 | 0.545 | 0.5887 | 0.068 |
| Initial learning | Random | 0.000118 | 0.000121 | eYFP - ChETA | -3.17e-06 | 1.91e-05 | 40 | -0.166 | 0.8690 | -0.021 |
| Initial learning | Regular | 0.000365 | 0.000371 | eYFP - ChETA | -6.30e-06 | 1.91e-05 | 40 | -0.330 | 0.7433 | -0.041 |
| Initial learning | Small-world | 0.000175 | 0.000178 | eYFP - ChETA | -3.29e-06 | 1.91e-05 | 40 | -0.172 | 0.8643 | -0.021 |
| Reversal learning | Goal-directed | 0.000372 | 0.000443 | eYFP - ChETA | -7.09e-05 | 2.83e-05 | 61 | -2.503 | 0.0150 | -0.403 |
| Reversal learning | Random | 0.000118 | 0.000118 | eYFP - ChETA | 3.40e-07 | 2.83e-05 | 61 | 0.012 | 0.9905 | 0.002 |
| Reversal learning | Regular | 0.000330 | 0.000359 | eYFP - ChETA | -2.92e-05 | 2.83e-05 | 61 | -1.030 | 0.3070 | -0.166 |
| Reversal learning | Small-world | 0.000173 | 0.000178 | eYFP - ChETA | -4.80e-06 | 2.83e-05 | 61 | -0.169 | 0.8661 | -0.027 |
